# Vascular-Perfusable Human 3D Brain-on-Chip

**DOI:** 10.1101/2025.09.18.676925

**Authors:** Alice E. Stanton, Rebecca L. Pinals, Aaron Choi, Nhat Truong, Eulim Kang, Alan Jiang, Claudia F. Lozano Cruz, Sydney Hawkins, Ryana Sarcar, Alexandra Volkova, Oisín King, Emre Agbas, Masayuki Nakano, Ching-Chi Chiu, Adele Bubnys, Shannon Wright, Colin Staab, Rami Bikdash, Emily Forden, Robert Langer, Li-Huei Tsai

## Abstract

Development and delivery of treatments for neurological diseases are limited by the tight and selective human blood–brain barrier (BBB). Although animal models have been important research and preclinical tools, the rodent BBB exhibits species differences and fails to capture the complexity of human genetics. Microphysiological systems incorporating human-derived cells hold great potential for modeling disease and therapeutic development, with advantages in screening throughput, real-time monitoring, and tunable genetic backgrounds when combined with induced pluripotent stem cell (iPSC) technology. Existing 3D BBB-on-chip systems have incorporated iPSC-derived endothelial cells but not the other major brain cell types from iPSCs, each of which contributes to brain physiology and disease. Here we developed a 3D Brain-Chip system incorporating endothelial cells, pericytes, astrocytes, neurons, microglia, and oligodendroglia from iPSCs. To enable this multicellular 3D co-culture in-chip, we designed a GelChip microfluidic platform using a 3D printing-based approach and dextran-based engineered hydrogel. Leveraging this platform, we co-cultured and characterized iPSC-derived brain-on-chips and modeled the brain microvasculature of *APOE4*, the strongest known genetic risk factor for sporadic Alzheimer’s disease. These 3D brain-on-chips provide a versatile system to assess BBB vascular morphology and function, investigate downstream neurological effects in disease, and screen therapeutics to optimize delivery to the brain.

**Significance Statement:** The blood–brain barrier (BBB) is both a contributing factor to neurological disease and a major obstacle to its treatment, yet human-relevant models remain limited. Most existing brain-on-chip systems incorporate only subsets of BBB cell types and cannot capture the full cellular complexity of the human neurovascular unit. Here, we establish a vascular-perfusable 3D Brain-Chip using human induced pluripotent stem cell-derived brain cells including endothelial cells, pericytes, astrocytes, neurons, microglia, and oligodendroglia. This system enables systematic analysis of human genetic risk factors, such as *APOE4* in Alzheimer’s disease, and provides a powerful platform to investigate BBB function and dysfunction and accelerate the development of more effective neurological therapies.

## Introduction

Neurological conditions are the leading cause of illness worldwide^1^, though the development of effective disease-modifying therapies for CNS disorders has been notoriously difficult and failure-prone.^2^ Contributing to this high failure rate are gaps in understanding of human disease mechanisms, limited technologies to study them, and the restrictive human blood–brain barrier (BBB), which most compounds fail to cross.^3^ The BBB is composed of lumenized networks of brain microvascular endothelial cells (BMECs) linked by tight junctions to form capillaries, the smallest blood vessels, which average 7–9 μm in diameter.^4^ These narrow dimensions dictate the distribution of blood cells, resistance to flow, and transport properties across the barrier.^5^ On the abluminal surface, BMECs are ensheathed by basement membrane proteins^6^ and pericytes^7^ and then astrocytic endfeet.^8^ The development and maintenance of the BBB are largely controlled by the other cell types comprising the neurovascular unit (NVU): pericytes, astrocytes, neurons, microglia, and oligodendroglia.^8^ Neuronal interactions with the endothelium are thought to contribute to barrier properties.^9^ While less is known about the effect of other glial cell types on barrier function and capillary phenotypes, recent studies indicate that microglia associate with capillary vessels, potentially influencing vascular tone,^10^ as do oligodendroglia, which appear to directly contact the vascular basement membrane.^11^

Humans differ from other species in BBB transcriptional signatures,^12^ permeability and transport properties,^13^ and even cellular components.^14,15^ *In vitro* brain models enable studies in human cell-based contexts, holding promise for mimicking aspects of human physiology and dissecting disease mechanisms, given the unique human genetic and gene regulatory mechanisms implicated in neurological diseases and the ability to construct such models from patient-specific cells.^16^ Simple two-dimensional (2D) cultures of BMEC monolayers alone or in co-culture with some number of other BBB cell types have been constructed as a research tool, such as in Transwell systems^17^ or multi-compartment microfluidic chips.^18^ However, 2D systems show lower barrier function, reduced tight junction expression, and transcriptional profiles distinct from three-dimensional (3D) systems.^19^ These cultures can be derived from primary or immortalized human cells,^18^ though their resultant barrier function has been less than physiological^19^ and tissue availability and reproducibility impose limitations. A promising cell source is human induced pluripotent stem cells (iPSCs), which enable modeling the BBB in a patient-specific and genotype-specific manner, including with isogenic sets of cell lines to isolate the contribution of specific genetic variants.^16^ Attention must be paid to the specifics of the differentiation protocol to ensure the specification of the endothelial and not epithelial lineage.^20^

Self-assembled vascular-perfusable 3D neurovascular models have been previously reconstructed with a subset of brain cell types.^21–24^ This includes an endothelial cell-neuron co-culture with human iPSC-derived endothelial cells and human embryonic stem cell-derived motor neuron spheroids co-embedded in a collagen gel in-chip.^21^ In a triculture model of the BBB, iPSC-derived endothelial cells and human primary pericytes and astrocytes were co-seeded in fibrin in commercially available microfluidic devices.^22^ Building on these systems, chip platforms have been constructed with a vascular bed of primary human microvascular endothelial cells, pericytes, and astrocytes embedded in fibrin, surrounding a central neurosphere compartment with ReN cell (immortalized human neural progenitors) neurospheres surrounded by Matrigel.^23,24^

Genetic variants have been identified for many neurological diseases, including many putatively linked to BBB dysfunction. This provides mechanistic clues to potential intervenable mechanisms and opens the possibility of personalized treatments based on genetic signatures.^25^ Accordingly, the construction of human cell-based models from patient-specific iPSCs is particularly important. Further, different genetic variants have distinct functional consequences, the various cell types in the brain express mutant proteins to varying extents, and these factors can have cascading effects through inherent *in vivo* multicellular crosstalk.^26^ Mounting evidence suggests that glial cell types play key roles in physiological functions and a broad array of neurological diseases.^27–30^ To address gaps in understanding related to current frontiers, such as neuroinflammatory mechanisms,^28^ glial crosstalk with neuronal phenotypes,^29^ and myelination homeostasis and remyelination,^30^ all of which involve interactions with the BBB, it is critical to develop human cell-based systems in which the unique contributions of all relevant cell types are included.

Towards addressing these gaps, we have recently established the human multicellular brain (miBrain) model composed of all six major brain cell types, each derived from iPSCs, in a neuromatrix hydrogel scaffold consisting of polysaccharide dextran, proteoglycans, and basement-membrane mimics to form functional 3D immuno-glial-neurovasculature.^31^ However, to more comprehensively assess vascular phenotypes, it is important to construct a vascular-perfusable neurovascular unit. To construct a 3D microvascular-perfusable human iPSC-based multicellular Brain-Chip, we developed a microfluidic platform, GelChip, fabricated using a 3D printing-based device fabrication approach. We identified device dimensions that support the self-assembly of contiguous vessel networks in co-culture. We optimized the method for co-culturing human iPSC-derived BMECs, pericytes, astrocytes, neurons, microglia, and oligodendroglia along with human fibroblasts, and selected subsets of these cell types, and characterized the resultant vessel network properties and permeabilities. Finally, to model the effect of *APOE4*, the strongest genetic risk factor for late onset Alzheimer’s Disease, on the BBB, we construct co-cultures of *APOE4* compared to isogenic *APOE3* controls and characterize vascular phenotypes. This work thus forms the foundation for studies of cerebrovascular interactions with neuronal processes and of BBB properties in disease-specific contexts.

## Results

### Development of a microfluidic device amenable to complex 3D co-cultures within engineered hydrogels

Few examples exist for stable co-cultures of four or more cell types within a single compartment, given the complexity of satisfying microenvironmental factors for all cells simultaneously.^32^ Similarly, there are only a few pioneering demonstrations of perfusable 3D self-assembled microvasculature with iPSC-derived BMECs.^21,22^ We have recently demonstrated the self-assembly of 3D microvasculature with six iPSC-derived cell types in the miBrain platform.^31^ To construct a 3D vascular-perfusable miBrain, we sought to combine the six iPSC-derived brain cell types: BMECs, pericytes, astrocytes, neurons, microglia, and oligodendroglia, with human fibroblasts, encapsulating in neuromatrix hydrogel^31^ and seeding in a microfluidic platform (**Fig. 1A**). In commercially available microfluidic devices, reduced co-cultures of BMECs with fibroblasts in fibrin protein-based gels robustly form networks. In contrast, miBrain co-cultures fail to form contiguous vessel networks in fibrin (**Fig. 1B**). Similarly, poor network formation is observed for BMEC-fibroblast co-cultures in the engineered neuromatrix hydrogel and for miBrain cell types co-cultured in fibrin (**Fig. 1B**).

**Figure 1.**
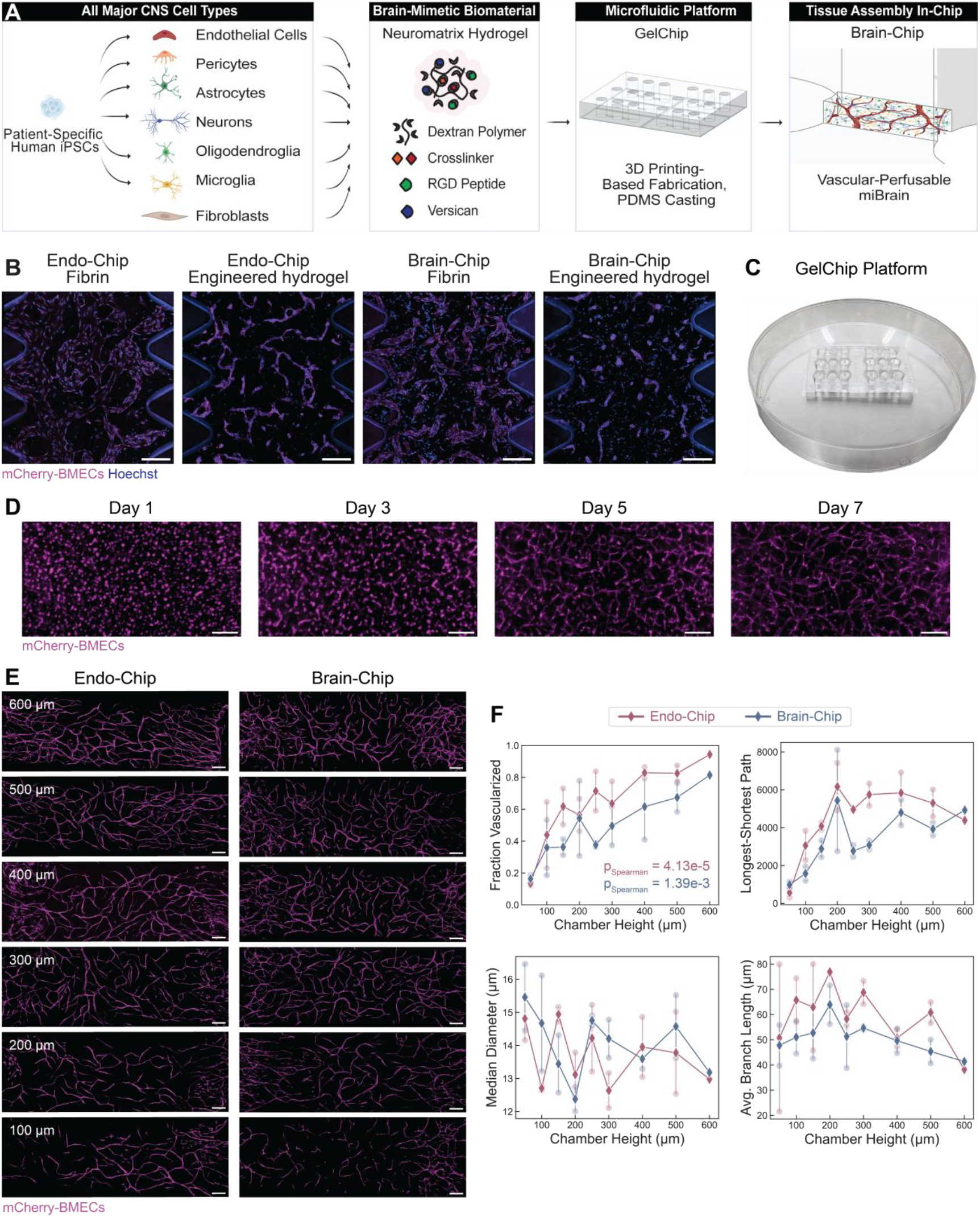
Engineering GelChip for culture of miBrain in microfluidic platform. (**A**) Schematic of Brain-Chip formation incorporating human iPSC-derived BMECs, pericytes, astrocytes, neurons, oligodendroglia, and microglia, co-encapsulated with fibroblasts in neuromatrix hydrogel and seeded into the microfluidic GelChip device chambers. (**B**) iPSC-derived BMECs cultured in AIM Biotech commercially available microfluidic devices in (*left to right*) fibrin with fibroblasts, neuromatrix hydrogel with fibroblasts, fibrin with all Brain-Chip cell types, and neuromatrix hydrogel with all Brain-Chip cell types (magenta: mCherry-BMECs; scale bars, 200 µm). (**C**) Macroscopic image of the GelChip. (**D**) BMECs in Endo-Chip at days 1, 3, 5, and 7 forming vascular networks (magenta: mCherry-BMECs; scale bars, 200 µm). (**E**) Optimization of GelChip chamber height for contiguous vessel network formation in-chip, displaying cultures in chips of (*top to bottom*) 600, 500, 400, 300, 200, and 100 µm height for (*left*) Endo-Chips of BMEC and fibroblast co-culture and (*right*) Brain-Chip of full miBrain and fibroblast co-culture (magenta: mCherry-BMECs; scale bars, 200 µm). (**F**) Quantification of vessel network parameters, as described in **Fig. S7**, for the fraction vascularized, longest-shortest path, median vessel diameter, and average branch length (red: Endo-Chip, blue: Brain-Chip; n=2 chips per group; individual replicates shown as semi-transparent circular markers, group means as solid diamond markers).

This result is anticipated given that engineered hydrogels, such as that which we developed and utilized for the miBrain, harbor innate swelling properties that, in confined spaces such as in microfluidic devices, can lead to mechanical forces on the cells detrimental to vessel lumen formation and network connectivity.^33^ Moreover, fibrin is not supportive of more complex, neuron-containing co-cultures such as the full miBrain.

We hypothesized that increasing the chamber dimensions could enhance 3D cell culture phenotypes (**Fig. S1A**). We devised an approach to fabricate devices of larger chamber dimensions via 3D printing molds for casting devices (**Fig. S1B**). We began with devices of 600 µm chamber height, 20 times larger than an average cell, which can be approximated as ∼30 µm, and 2.4 times larger than the commercially available microfluidic devices tested. Entry and exit ports flank either side of the chamber. In this GelChip microfluidic platform (**Fig. 1C**), BMECs co-cultured with fibroblasts in the engineered neuromatrix hydrogel formed networks within 5–7 days (**Fig. 1D**). Cell media (**Fig. S2, Table S1**) and cell compositional ratios (**Fig. S3**) were optimized to support vascular network formation. At 600 µm, Brain-Chips display robust protein immunoreactivity to canonical markers for vessels (**Fig. S4A**) with evident tight junctions (**Fig. S4B**), GFAP-positive astrocytes (**Fig. S4C**), neurons (**Fig. S4D**), and iPSC-derived microglia (iMG) (**Fig. S4E**). Thus, GelChips are supportive of iPSC-derived vessel networks, even within an engineered hydrogel and with higher complexity brain cell co-cultures.

We next tested the effect of chamber height on vessel network properties between 100 and 600 µm in 100 µm increments for co-cultures of miBrain cell types with fibroblasts and for reduced co-cultures of BMECs with fibroblasts, both in the engineered neuromatrix hydrogel (**Fig. 1E,F**). With increasing chamber height within this range, we observed increasing connectivity, quantified as the fraction vascularized. The longest–shortest path (defined as the maximum geodesic distance within the largest connected component) increases with chamber height up to ∼200 µm, at which point it plateaus at ∼3.5 mm, approximately equal to the full chamber length of 3.350 mm. This indicates that by 200 µm height, vessels form some paths spanning the chamber. However, the fraction vascularized continues to rise with greater chamber heights up to 600 µm, reflecting an expansion of contiguous vessel networks. Notably, increasing the overall chamber length further extended the longest–shortest path (**Fig. S5A**) and changing the width did not modify fraction vascularized (**Fig. S5B**). The median vessel diameter and average branch length do not change appreciably with chamber height, as anticipated and required to retain microvascular network formation capability. Therefore, we have established a platform for the culture of contiguous iPSC-derived vessel networks within engineered hydrogels in-chip, including in co-culture with the other five major brain cell types and fibroblasts.

### Microvascular networks within multicellular co-cultures in-chip

Seminal 3D printing-based methods were previously developed to construct lumenized, perfusable vessel networks of 150–800 µm diameter.^34^ Subsequent studies have demonstrated perfusion in 3D vessels down to 80–200 µm diameter, using a hybrid approach that combines bioprinting and biofabrication methods.^35^ Casting approaches in which the hydrogel is polymerized around a removable wire prior to cell seeding have led to perfusable 3D vessels of 150 µm diameter.^36^ However, human brain capillary vessels are thinner, roughly 7–9 µm diameter.^4^ In contrast, the thinnest perfusable 3D engineered vessels that have been demonstrated, to our knowledge, are roughly 40 µm in diameter,^21,24^ and are a product of cell self-assembly approaches.

Within this context, we characterized the vessels in simplified BMEC and fibroblast cultures in engineered neuromatrix hydrogel seeded in the GelChip microfluidic platform (Endo-Chip; **Fig. 2A**). We also tested reduced co-cultures of iPSC-derived BMECs and neurons with human fibroblasts (NVU-Chip; **Fig. 2B**) and iPSC-derived BMECs, pericytes, and astrocytes with human fibroblasts (BBB-Chip; **Fig. 2C**), in addition to all of the miBrain cells (iPSC-derived BMECs, pericytes, astrocytes, neurons, oligodendroglia, and microglia) with human fibroblasts (Brain-Chip; **Fig. 2D**). Representative images from day 7 are included in **Fig. 2**, with an extended time course in **Fig. S6**. Vessels were analyzed using a supervised machine-learning based approach (**Fig. S7**) as detailed in the Methods section. The vessel networks increased in network connectivity over time in culture, with Endo-Chips and NVU-Chips plateauing in the fraction vascularized sooner (by day 5) and attaining higher overall values with less variance compared to the BBB-Chip and Brain-Chip (**Fig. 2E**). BBB-Chips were also more similar to Brain-Chips in the Euler characteristic (**Fig. 2F**), a topological invariant representing network connectivity, with lower values indicating greater vascular network interconnectedness. At day 7, the average vessel diameter across all co-cultures is 13.6 μm (interquartile range, 10.6-15.7 μm; **Fig. 2G**), and the average branch length was 51.7 μm (interquartile range, 21.5-71.0 μm; **Fig. 2H**). The distribution of vessel diameters in our engineered co-cultures are higher than those reported for human *in vivo* capillaries^37^ (**Fig. 2G**), but an improvement compared to all previous attempts that we are aware of, with BBB-Chips and Brain-Chips being particularly close to the *in vivo* condition. Further, the distribution of branch lengths across co-cultures tracks very similarly to *in vivo* vessels (**Fig. 2H**). Overall, we found that our co-cultures produced with vessel diameters and branch lengths more similar to those found *in vivo*, and that the cell type composition of the co-cultures influences these network parameters.

**Figure 2.**
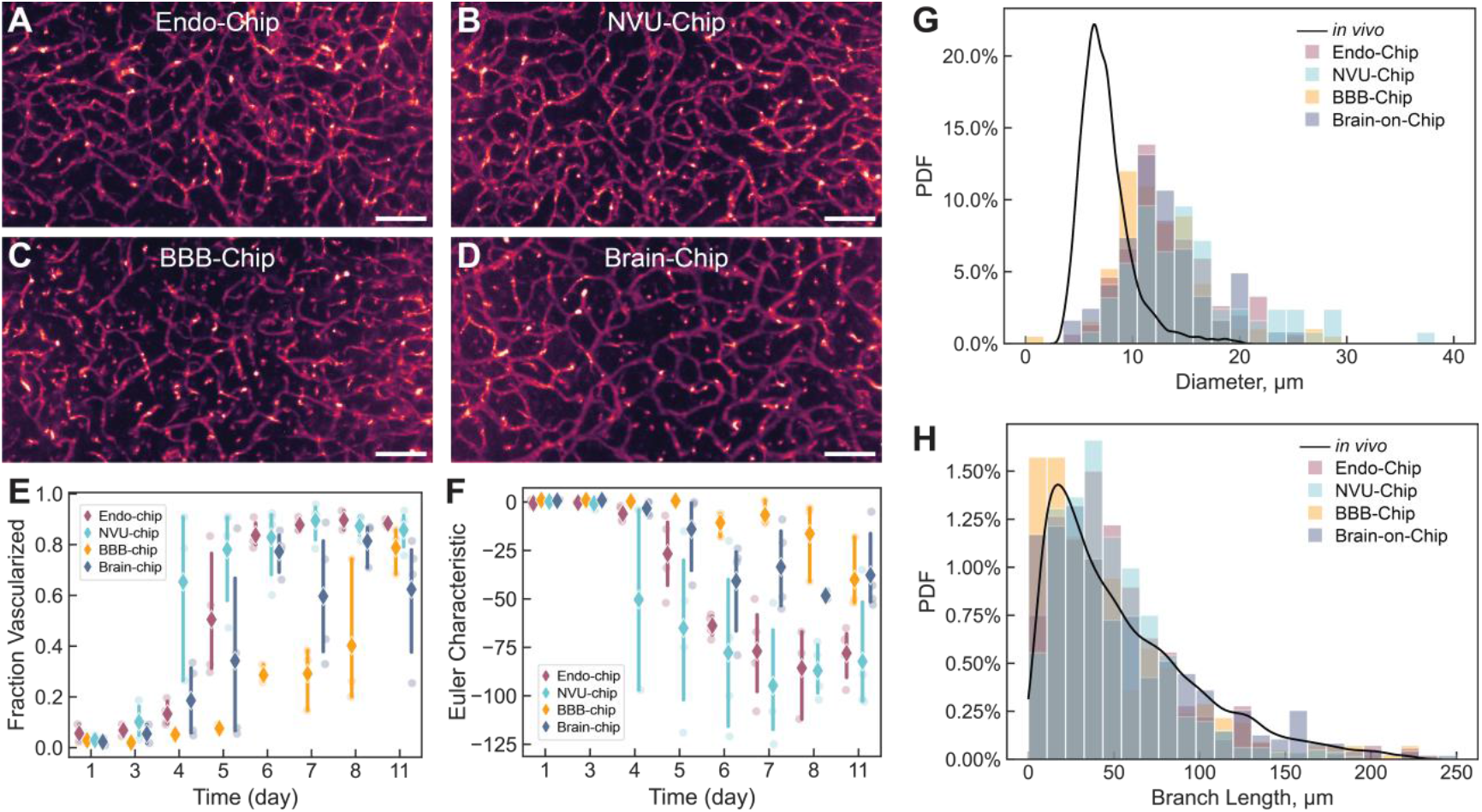
Vessel networks across co-culture compositions. Local thickness mapping of vessel networks in (**A**) Endo-Chips of BMEC-fibroblast co-culture, (**B**) NVU-Chips of neuron, BMEC, fibroblast co-culture, (**C**) BBB-Chips of astrocyte, pericyte, BMEC, fibroblast co-culture, and (**D**) Brain-Chips of full miBrain (neuron, microglia, oligodendroglia, astrocyte, pericyte, BMEC)-fibroblast co-culture (day 7; scale bars, 200 µm). (**E-F**) Quantification of vessel networks across co-culture compositions (red: Endo-Chip; light blue: NVU-Chip; yellow: BBB-Chip; violet: Brain-Chip) for (**E**) fraction vascularized and (**F**) Euler characteristic (n=3-4 chips per group; individual replicates shown as semi-transparent circular markers, group means as solid diamond markers). Distribution of (**G**) vessel diameters and (**H**) vessel branch lengths in comparison to human cerebral cortex capillaries (black line; Cassot, F. *et al* Microcirculation 2006).

### Perfusion of microvascular networks in-chip

To establish perfusion of solutes through the microvasculature lumen, we employed a method to connect the 3D vessels in co-cultures with the microfluidic chip inlet and outlet ports. We first microdissected excess hydrogel at the chamber edges to render the hydrogel flush with the chamber and improve access to vessels within the chamber (**Fig. S8**). Additional BMECs were then seeded into the ports to anastomose with the 3D vessels inside the chamber, using a previously reported method.^38^ Using this approach in a proof-of-principle experiment, Endo-Chips containing larger-diameter GFP-HUVEC (rather than BMEC) vessels were perfusable with 1 µm microparticles (**Fig. 3A**). We established that Brain-Chips containing smaller-diameter iPSC-BMEC microvessels were perfusable with small-molecule tracers (**Fig. 3B**), with perfusion through vessels as narrow as ∼16 µm diameter.

**Figure 3.**
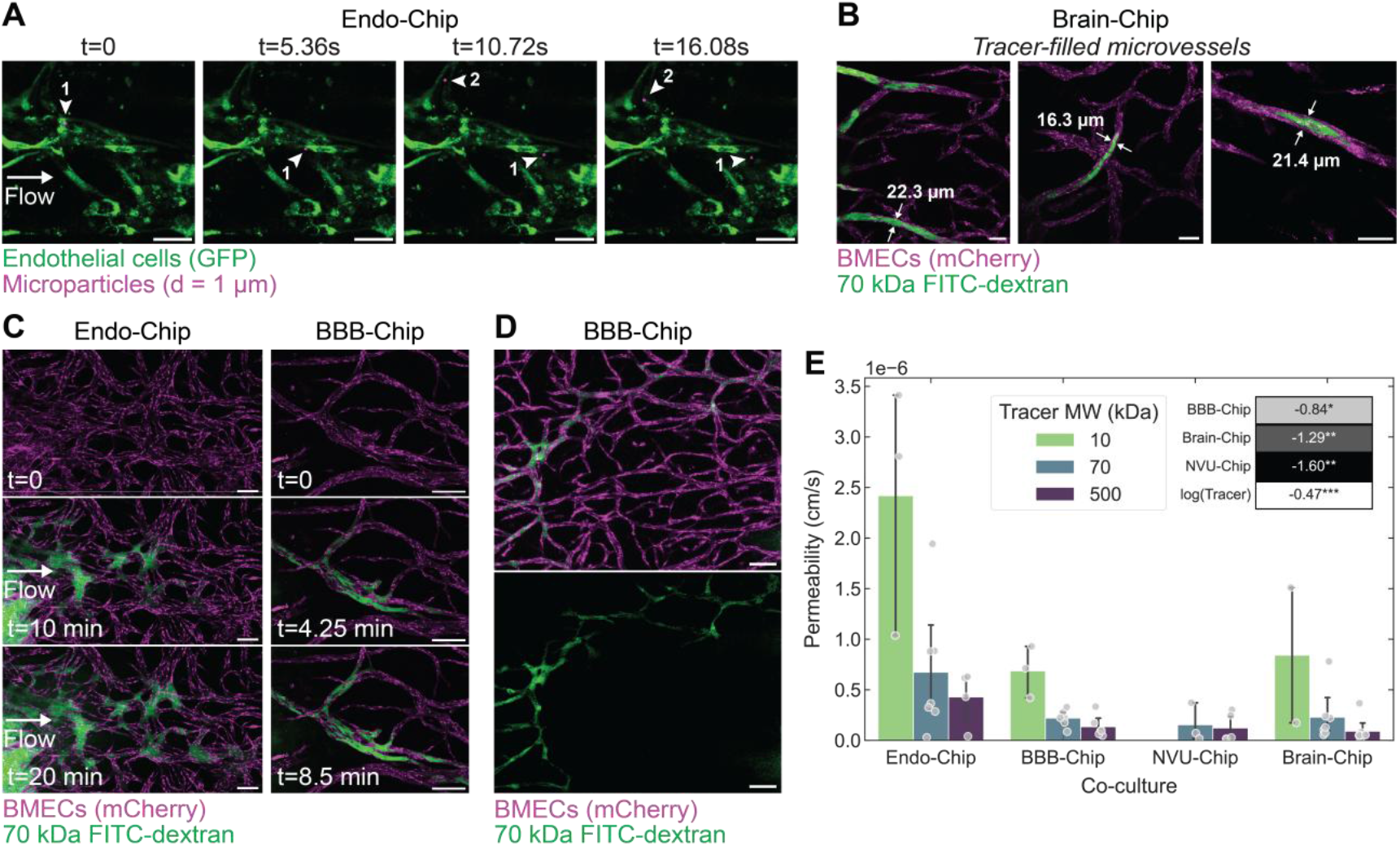
Luminal perfusion of vessel networks in GelChip. (**A**) Perfusion of Endo-Chip 3D vessels with 1 µm particles (green: GFP-HUVECs, magenta: PEGylated polystyrene microparticles; scale bars, 50 µm). (**B**) Examples of perfusion through microvasculature within Brain-Chip 3D vessels with 70 kDa FITC-dextran (magenta: mCherry-BMECs, green: FITC-dextran; scale bars, 50 µm). (**C**) Examples of perfusion of 70 kDa FITC-dextran through (*left*) Endo-Chip and (*right*) BBB-Chip (*top to bottom*) over time (magenta: mCherry-BMECs; green: FITC-dextran; scale bars, 100 µm) (**D**) Post-perfusion image of BBB-Chip (magenta: mCherry-BMECs; green: FITC-dextran; scale bars, 100 µm). (**E**) Quantification of permeability across vascular barrier for Endo-Chip, BBB-Chip, NVU-Chip, and Brain-Chip across molecular sizes of 10, 70, and 500 kDa FITC-dextran tracers (n=2-8 chips per group, with individual chips shown as markers). Inset: regression coefficients from log-transformed permeability values, showing relative differences in barrier tightness across co-culture types relative to Endo-Chip reference and inverse dependence on tracer molecular weight.

We used a panel of FITC-dextran tracers to visualize perfusion, as shown in representative images for Endo-Chip and BBB-Chip cultures (**Fig. 3C,D**), and to quantify the permeability coefficients across vessels within all co-cultures (**Fig. 3E**). As expected, permeability increased with decreasing tracer size, from 500 to 70 to 10 kDa. To interpret the trends, we performed a multivariable linear regression of log-permeability against culture type (relative to Endo-Chips) and log-tracer mass (**Fig. 3E** inset). The tracer regression coefficient of −0.47 closely matches the classical scaling of dextran diffusivity as a polymer in bulk solution (*D*∝ *M*^-0.5^),^39^ consistent with size-dependent transport restriction. Moreover, this analysis demonstrates that adding supporting brain cell types progressively tightened barrier function, wherein the lowest permeability was observed with the inclusion of neurons in the NVU-Chips and Brain-Chips.

### Genotype-specific modeling with Brain-Chip

Constructing these cultures from iPSC-derived cells enables the examination of vessel properties from different human individuals and from different genotypic backgrounds. As one proof-of-concept, we modeled the vascular phenotypes in *APOE4* cultures, the genotype most strongly associated with late-onset Alzheimer’s Disease, using an isogenic set of iPSCs, *APOE3* parental and its pair CRISPR-edited to *APOE4*. Studies have indicated that *APOE4* affects the BBB capillary basement membrane area^40^ and BBB breakdown in humans^41–43^ and BBB dysfunction in *APOE4* transgenic mouse models.^44–48^

BBB-Chips of *APOE4* compared to *APOE3* background had altered vascular network properties, with *APOE4* exhibiting less network connectivity, as evidenced by decreased percent coverage and fraction vascularized metrics. The Euler characteristic, indicative of network topological complexity, was also lower in *APOE4* BBB-Chips (**Fig. 4A,B**). Concomitantly, tight junction barrier function was lower in *APOE4* vasculature compared to that of *APOE3* in Brain-Chips for ZO-1 immunoreactivity (**Fig. 4C**).

**Figure 4.**
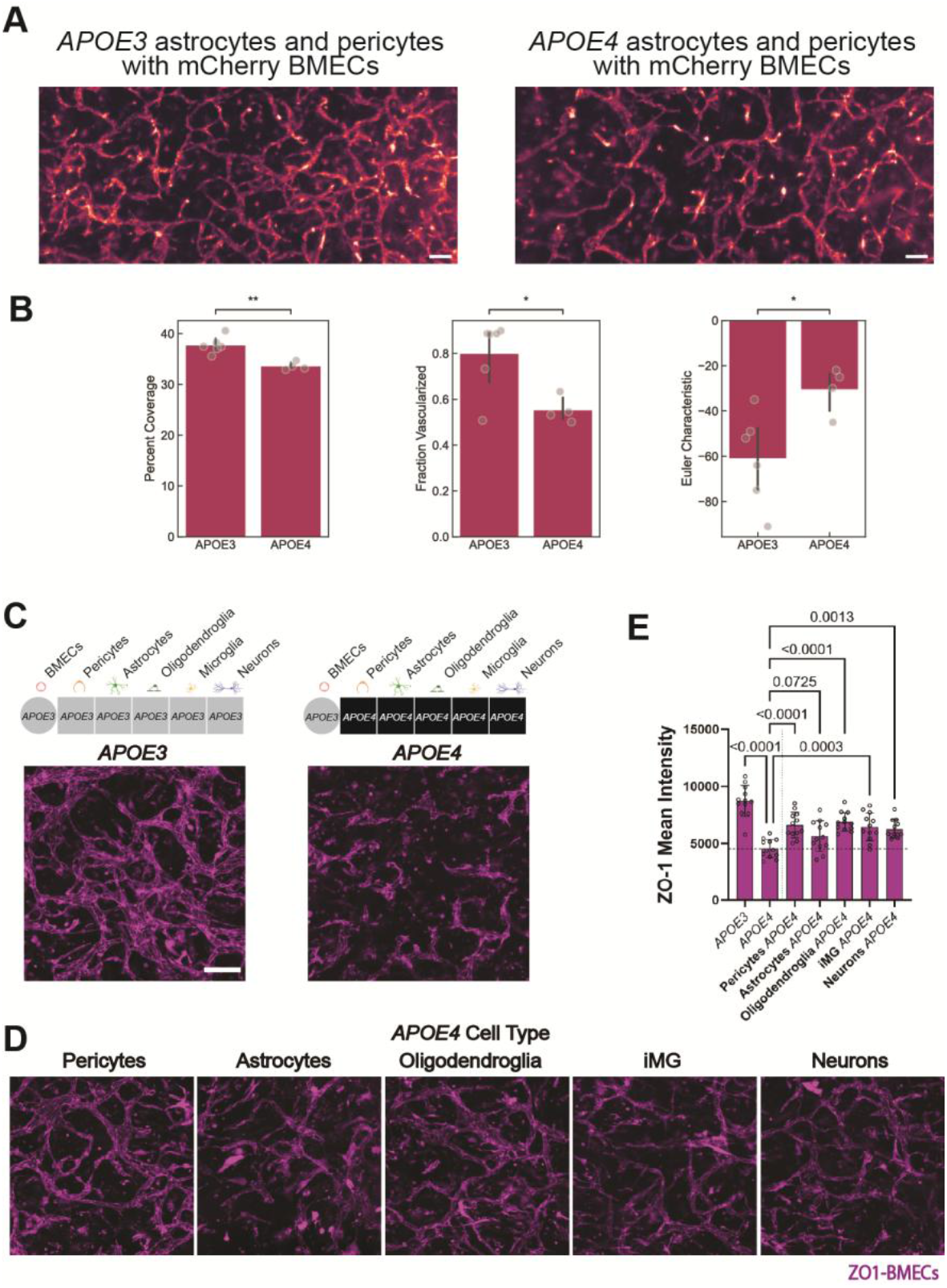
Modeling *APOE4-*associated vascular defects. (**A**) mCherry BMEC vessels with *APOE3* versus *APOE4* pericytes and astrocytes in BBB-Chips (magenta: mCherry BMECs; scale bar, 100 µm), with (**B**) complementary characterization of vessel properties. (**C**) ZO-1-BMECs in *APOE3* versus *APOE4* Brain-Chips and (**D**) *APOE3* miBrains with one cell type of *APOE4* background: (*left to right*) pericytes, astrocytes, oligodendroglia, microglia, or neurons (magenta: ZO-1-BMECs; scale bar, 100 µm). (**E**) Quantification of tight junction mean intensity across conditions (n=4 replicates with n=12 FOVs/ group).

Harnessing the modularity of the system to test contributions of individual cell types to this dysfunction, in parallel, we cultured *APOE3* Brain-Chips in which one cell type was replaced by its *APOE4* cell counterpart (**Fig. 4D**). Interestingly, Brain-Chips with *APOE4* astrocytes had markedly decreased barrier function (**Fig. 4E**). This decreased ZO-1 immunoreactivity was not as low as *APOE4* Brain-Chips cultured in parallel but was the most disruptive of the individual cell types. Thus, the Brain-Chips can be used as a highly modular investigative tool to research and recapitulate key clinical observations, including in the contexts of disease and genetic risk.

## Discussion

Human cell-based models offer new possibilities of utility for mechanistic inquiry, therapeutic discovery, and drug development. Here we have developed a microfluidic platform supportive of perfusable vessel networks within 3D co-cultures, including in cultures using engineered neuromatrix hydrogels as the cell scaffolding material. Given the growing desire to move away from animal-derived proteins in such *in vitro* systems and to identify suitable hydrogel systems that can better mimic aspects of the native human tissue, the compatibility of this platform with engineered neuromatrix hydrogel is significant. Importantly, this chemically defined extracellular matrix supports both robust vascular network formation and integration of diverse brain cell types, overcoming limitations of other hydrogel matrices. The innate swelling property of hydrogels has been a critical bottleneck inhibiting their use in confined microfluidic environments. We identified the microfluidic chamber height to be a key parameter in modulating 3D cell network phenotypes in these cultures. While the 600 µm height led to perfusable vessels in the Endo-Chip, BBB-Chip, NVU-Chip, and Brain-Chip co-cultures, it is possible that alternative cell type combinations would benefit from testing a range of chamber heights.

To facilitate 3D vessel perfusion within our co-cultures, we have further found that microdissection of the outer perimeter of 3D cultures in the engineered hydrogel, prior to interface seeding with endothelial cells, to be critical to enabling perfusion. Here we used simple dissection forceps to remove this outer perimeter, though future work could develop tools to further automate and standardize this process.

Leveraging this platform, we have demonstrated successful 3D vessel network formation and perfusion within Endo-Chip, BBB-Chip, NVU-Chip, and Brain-Chip co-cultures. Notably, the 3D vessels contained therein are ∼10-15 µm in diameter, closely approaching that of human capillary dimensions, and significantly smaller than those reported in prior engineered vessel models. The branch length distribution mirrors that of the human BBB microvasculature, underscoring the ability of this system to recapitulate native-like vascular architecture. Permeability measurements provide additional confirmation of barrier function, with BBB-Chips showing an average permeability of 2.20 × 10^−7^ cm/s for 70 kDa FITC-dextran, comparable to reported values in rat BBB,^49,50^ and NVU-Chips averaging 1.54 × 10^−7^ cm/s, similar to prior 3D endothelial cell-astrocyte-neuron co-cultures in microfluidic chips.^51^

Harnessing iPSC-derived cell types and developing a Brain-Chip inclusive of all six major brain cell types holds new possibilities for the construction of disease models of patient-specific and genotype-specific backgrounds. Here we investigated the effect of *APOE4* cell types using one set of iPSC lines, which should be extended to multiple cell lines in the future to corroborate findings. Vascular network analysis could be further applied to these cell type combinations to further dissect cell type-specific effects. While we used human primary fibroblasts, future work could integrate iPSC-derived fibroblasts for fully isogenic comparisons. Additional work could also further optimize the success rate for perfusion, as we observed with increasing number of distinct cell types, decreased odds of vessel entry points at the chip inlets, which are needed for perfusion. Integration of liquid handling and robotics that increase precision in cell counts and seeding procedures could be advantageous in increasing homogeneity in co-culture setup and addressing this limitation.

In sum, we have developed a Brain-Chip platform with notably enhanced features in terms of cell type complexity and vessel network properties that could be harnessed for assessment of BBB transport and permeability, including in disease contexts. At the same time, we have developed a microfluidic device that could hold new possibilities for microphysiological systems with engineered hydrogels and complex co-cultures.

## Materials and Methods

Detailed information for all experimental methods and materials is provided in SI Appendix.

### Cell lines

Human iPSC lines used in this study were previously described.^31^ In brief, an endogenously expressing mCherry iPSC line was used predominantly for BMECs (parental line CS-0009-01; from a healthy *APOE2/3* male donor of unknown age). A ZO1-mEGFP line (Allen Institute for Cell Science parental line WTC-11, Coriell GM25256; from a healthy 30-year-old male donor) was also used for BMECs. An *APOE3/3-*parental iPSC line (AG09173; from a healthy 75-year-old female donor) was predominantly used for all other cell types in Endo-Chip, BBB-Chip, NVU-Chip, and Brain-Chip cultures. Comparisons to *APOE4* were conducted with an isogenic iPSC line that was previously CRISPR-edited to *APOE4/4* (AG09173-1336) by the Picower Institute for Learning and Memory iPSC Facility.^52^ Normal human lung fibroblasts were obtained from Lonza (CC-2512). GFP-expressing human umbilical vein endothelial cells (GFP-HUVECs) were obtained from Angio-Proteomie (cAP-0001GFP).

### Microfluidic GelChip device fabrication

Microfluidic devices were designed with varying chamber dimensions and with inlets and outlets directly accessible to the cell culture chamber (AutoCAD). The negative of these designs were 3D printed with grey resin (Formlabs) using fine layer thickness in an SLA 3D Printer (Formlabs). Resulting molds were washed vigorously in an automated washing unit (Formlabs) with 90% IPA, cured in a UV oven (Formlabs), and coated by depositing a thin 2 µm layer of parylene to prevent unreacted monomer leaching from inhibiting downstream polymer crosslinking. Devices were cast by pouring vigorously mixed 1:10 crosslinker-to-base PDMS (Krayden Dow) on molds, degassing for 45 minutes, and curing at 80° C overnight. The following day, devices were removed from molds and bonded to thin glass coverslips (Thickness #1; Ted Pella) using a plasma cleaner (Harrick Plasma). Devices were baked for another hour at 80° C prior to autoclaving and cell culture. GelChips were also fabricated by a contractor for some cultures using their proprietary method to fabricate devices in PMMA (Parallel Fluidics). Cultures were also tested in commercially available microfluidic devices (AIM Biotech).

### Cell culture

iPSCs were cultured using the mTeSR1 medium system (STEMCELL Technologies, 85850) on 2% v/v hESC-qualified Matrigel (Corning, 354277) coated vessels. Cells were differentiated as previously described.^31^ Cells were dissociated with Accutase (STEMCELL Technologies, 07922) and encapsulated in either neuromatrix hydrogel (Sigma, TrueGel3D Hydrogel Kits with cell-degradable cross-linker TRUE7, with optimized inclusion of RGD and versican peptides) as previously described^31^ or fibrin hydrogel (3 mg/mL fibrinogen from Millipore Sigma, F8630; 2 U/mL thrombin from Millipore Sigma, T4648) as previously described.^53^ To seed microfluidic devices, hydrogel precursor solution with all cells in suspension was injected into the cell culture chamber, with volumes of approximately 2-5 µL depending on device design. Neuromatrix hydrogels were polymerized for 25 min at 37°C, 5% CO_2_. Fibrin hydrogels were polymerized at room temperature for 15 min, with flipping of the chip approximately every 30 seconds.^54^ Media was composed of EGM-2 MV from Lonza (CC-3202) supplemented with 50 ng/mL VEGF-165 (human recombinant protein, PeproTech, 100-20) for one week, reduced to 10 ng/mL thereafter. BBB-Chip media additionally included 10 ng/mL of PDGF-BB (human recombinant protein, PeproTech, AF-100-14B) while NVU-Chip additionally included 1 nM of SAG (potent Smoothened receptor agonist, Tocris, 4366). Brain-Chip media was composed of EGM-2 MV with all supplements except FBS, in addition to VEGF-165 as described above, 1% v/v AGS (ScienCell, 1852), 2% v/v B-27 (Gibco, 17504044), 1 µM cAMP (N6,2′-O-Dibutyryladenosine 3′,5′-cyclic monophosphate sodium salt, Millipore Sigma, D0627), 10 ng/mL NT-3 (Peprotech, 450-03), 1 nM SAG (Tocris, 4366), and 25 ng/mL M-CSF (human recombinant protein, PeproTech, 300-25).

Endo-Chips were seeded with 5 × 10^6^ cells/mL endothelial cells (ECs) and 1.5 × 10^6^ cells/mL fibroblasts. BBB-Chips were seeded with 5 × 10^6^ cells/mL ECs, 0.3125 × 10^6^ cells/mL pericytes, 0.625 × 10^6^ cells/mL astrocytes, and 1.25 × 10^6^ cells/mL fibroblasts. NVU-Chips were seeded with 4 × 10^6^ cells/mL ECs, 3 × 10^6^ cells/mL neurons, and 1.5 × 10^6^ cells/mL fibroblasts. Brain-Chips were seeded with 5 × 10^6^ cells/mL ECs, 0.3125 × 10^6^ cells/mL pericytes, 0.625 × 10^6^ cells/mL astrocytes, 4 × 10^6^ cells/mL neurons, 0.3125 × 10^6^ cells/mL oligodendrocyte progenitor cells (OPCs), and 1.25 × 10^6^ cells/mL fibroblasts, and 0.3125 × 10^6^ cells/mL microglia.

Co-cultures were fed daily by medium exchange through the inlets and outlets. On days 4 then 5, inlets then outlets were respectively seeded with additional BMECs as previously reported,^38^ with the following modifications. First, the hydrogel perimeter at each port was microdissected with forceps (DR Instruments) to remove the acellular rim and enhance access to the 3D vessel networks within the hydrogel. Ports were coated for 1-2 h at 37°C, 5% CO_2_ with basement membrane proteins selected based on the device material and with sufficient volume to cover the lower surface. For in-house fabricated devices (glass bottom), ports were coated with 100 µL volume of 50 µg/mL fibronectin (Corning, 354008) diluted in DPBS. For contractor fabricated devices (PMMA bottom), ports were coated with 73 µL volume of 50 µg/mL collagen (Gibco, A1048301) diluted in DPBS. BMECs (0.35 × 10^6^ cells/mL) were seeded with the corresponding volumes (100 µL for in-house or 73 µL for contractor), devices were tilted ∼70° in the incubator for 2 h, and seeding was repeated until a uniform interface was observed (typically twice). Chips were then tilted ∼30° and incubated overnight.

## Acknowledgments

The authors acknowledge Dr. Siddarth Krishnan for assistance in coating device molds and the MIT Koch Institute Flow Cytometry Core at the Swanson Biotechnology Center for assistance with flow cytometry. The authors would like to acknowledge funding through National Institutes of Health grants 3-UG3-NS115064-01S1, 1F32AG072813-01, and 1K01AG083734-01, as well as generous support from the BT Charitable Foundation, the Freedom Together Foundation, The Picower Institute for Learning and Memory, the Robert A. and Renee E. Belfer Family, Eduardo Eurnekian, David B. Emmes, and Kathleen and Miguel Octavio. A.E. Stanton acknowledges support from F32AG072813, K01AG083734, and the Kavanaugh Fellowship. R.L. Pinals acknowledges support from the Schmidt Science Fellows program in partnership with the Rhodes Trust and the Burroughs Wellcome Fund Career Award at the Scientific Interface (CASI). This work was supported in part by the Koch Institute Support (core) Grant P30-CA14051 from the National Cancer Institute. We thank the Koch Institute’s Robert A. Swanson (1969) Biotechnology Center for support, specifically Peterson (1957) Nanotechnology Materials Core Facility (RRID:SCR_018674).

