## Supporting Information for "Vascular-Perfusable Human 3D Brain-on-Chip" (from another version of this preprint: version 1, 2025-09-19)

#### Supporting Information for
### Vascular-Perfusable Human 3D Brain-on-Chip

Alice E. Stanton[1*†], Rebecca L. Pinals[2,3*†], Aaron Choi[2,3], Nhat Truong[2,3], Eulim Kang[2,3], Alan Jiang[2,3], Claudia F. Lozano Cruz[2,3], Sydney Hawkins[1], Ryana Sarcar[1], Alexandra Volkova[1], Oisín King[2,3], Emre Agbas[2,3], Masayuki Nakano[2,3], Ching-Chi Chiu[2,3], Adele Bubnys[2,3], Shannon Wright[2,3], Colin Staab[2,3], Rami Bikdash[1], Emily Forden[1], Robert Langer[1,4,5,6,7*], Li-Huei Tsai[2,3,8*]

[1]Koch Institute, Massachusetts Institute of Technology; Cambridge, USA.
[2]Picower Institute for Learning and Memory, Massachusetts Institute of Technology; Cambridge, USA.
[3]Department of Brain and Cognitive Sciences, Massachusetts Institute of Technology; Cambridge, USA.
[4]Department of Chemical Engineering, Massachusetts Institute of Technology; Cambridge, USA.
[5]Department of Anesthesiology, Boston Children's Hospital; Cambridge, USA.
[6]Division of Health Science and Technology, Massachusetts Institute of Technology; Cambridge, USA.
[7]Institute for Medical Engineering and Science, Massachusetts Institute of Technology; Cambridge, USA.
[8]Broad Institute of Harvard and Massachusetts Institute of Technology; Cambridge, USA.

*Co-corresponding authors. Li-Huei Tsai, Robert Langer, Alice E. Stanton, Rebecca L. Pinals.

†These authors contributed equally to this work.

**This PDF file includes:**

Supplementary Methods
Figs. S1 to S8
Table S1
SI References

#### Supplementary Methods

**Microscopy.** Confocal microscopy imaging was performed on a Zeiss LSM900 confocal microscope with ZEN Blue software (v3.9) using 10×/0.3 NA and 20×/0.8 NA Plan-Apochromat objectives (Zeiss). Excitation was provided by 405, 488, 561, and 640 nm lasers on separate imaging tracks, with emission windows set to 410-470, 492-550, 564-625, and 650-700 nm, respectively. Laser-blocking filters were applied to all excitation lines and a 470 nm shortpass filter was used with the 405 nm laser. The pinhole was set to 1 Airy unit for each channel. Laser intensity and detector gain were adjusted to avoid saturation, then maintained uniformly across each sample set. Z-stacks were collected to capture the full sample thickness at Nyquist step sizes.

Live imaging was performed on the same LSM900 confocal microscope as described, with a 37 °C heated lid under humidified 5% $CO_2$ atmosphere, or on an Agilent BioTek Cytation C10 confocal imaging reader, with the chamber maintained at 37 °C and 5% $CO_2$. For the latter, images were acquired with Gen5 Image Prime software (v3.16) using a 20× Plan Fluorite objective (Olympus) and a 40 µm Nipkow spinning disk. Fluorescence was acquired with filter sets at 390/442 nm (DAPI), 472/520 nm (GFP), 556/600 nm (TRITC), and 635/685 nm (Cy5). Z-stacks were collected to capture the full sample thickness. Images for vascular network visualization as a function of time were also collected on the EVOS imaging system (Invitrogen).

**Vessel network analysis.** Vessel images were analyzed in Fiji (ImageJ distribution NIH, v2.14.0/1.54g) with custom macros. Maximum-intensity projections of the vessel channel (mCherry, 561 nm channel) were cropped to a consistent ROI, then batch-processed to contrast-normalize by linear rescaling (only for images collected on the EVOS imaging system), segmented with a pretrained Trainable WEKA Segmentation classifier to isolate vessel structures from background, converted to binary masks, despeckled and skeletonized. Mask-level morphology and local thickness were quantified with the BoneJ plugin, yielding outputs including area measurements, vessel morphology, local-thickness histograms, and Euler characteristic. Network-level metrics were obtained with the Analyze Skeleton (2D/3D) plugin, yielding outputs including skeleton branch information.

Vascular network features were quantified from segmented and skeletonized images. Fraction vascularized is defined as the proportion of total branches present in the largest connected component within the vascular network. The longest-shortest path is calculated as the length of the maximum geodesic distance (shortest path) within the largest connected component. Median diameter is calculated as the median vessel thickness from local-thickness histograms. Average branch length is calculated as the mean length of all skeleton branches. The Euler characteristic is calculated from the largest connected component (1) minus the number of holes. Percent coverage is defined as the ratio of vascular area to total image area.

**Vessel perfusion assays.** At day 7-9, vascular permeability was tested as previously described.[1] Briefly, cell culture medium was gently removed from device ports and replaced with 0.1 mg/mL FITC-dextran (Millipore Sigma) diluted in the same medium. Chips were live-imaged to capture both FITC and mCherry signal at two timepoints spaced approximately 3-5 min apart.

To quantify vascular permeability in-chip, paired z-stack images were acquired at two timepoints of tracer perfusion and processed in Fiji with a custom macro. Image stacks of the tracer (FITC-dextran of varying molecular weights, 488 nm channel) were cropped in xy, trimmed in z, background-subtracted, contrast-enhanced, and segmented using a pretrained Trainable Weka Segmentation 3D classifier to isolate vessel structures. Segmented volumes were imported into the 3D Manager plugin for geometric analysis and the 3D Intensity Measure plugin to extract intravascular versus extravascular fluorescence intensities. Permeability coefficients were calculated from intensity changes across the vessel boundary using paired pre- and post-perfusion datasets according to:

$$P = \frac{V_{\mathrm{m}}\, \Delta I_{\mathrm{m}}}{\Delta t\, A_{\mathrm{v}}(I_{\mathrm{v}} - I_{\mathrm{m}})}$$

where $V_m$ is the hydrogel matrix volume, $I_m$ and $I_v$ are fluorescence intensity in the vessel and hydrogel matrix at the initial time, respectively, $\Delta t$ is the time interval between collected timepoints, $A_v$ is the surface area of the vessel, and $\Delta I_m$ is the difference in fluorescence intensity in the hydrogel matrix between at the initial and final time points. This approach makes the following assumptions: (1) fluorescence intensity is proportional to tracer concentration, (2) hydrogel matrix is homogenous and uniformly permeable, (3) tracer concentration in the vessel is constant throughout the measurement, (4) tracer undergoes negligible photobleaching, and (5) no transport across device boundaries.

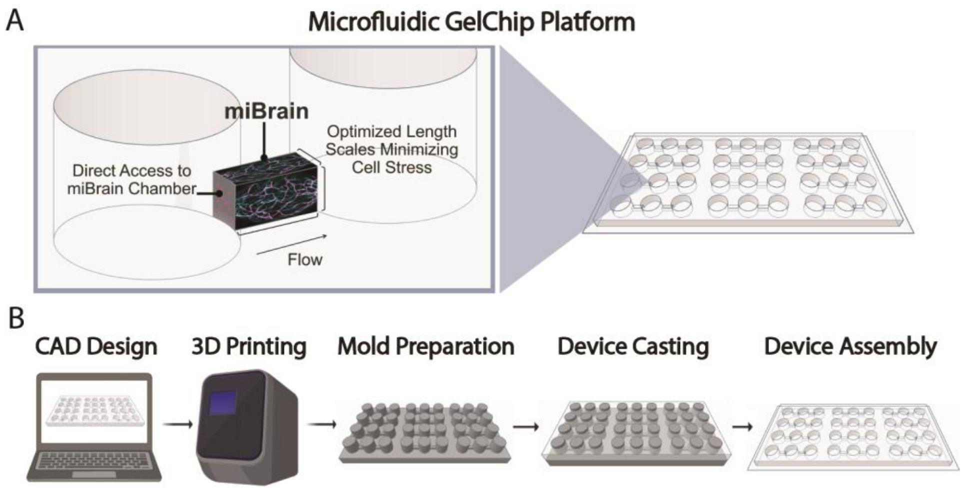


**Fig. S1. Microfluidic device 3D printing-based method.** (**A**) Schematic of GelChip platform and (**B**) method of 3D SLA printing molds, preparing molds with coating, casting PDMS on molds, and assembling devices by bonding to glass coverslips.

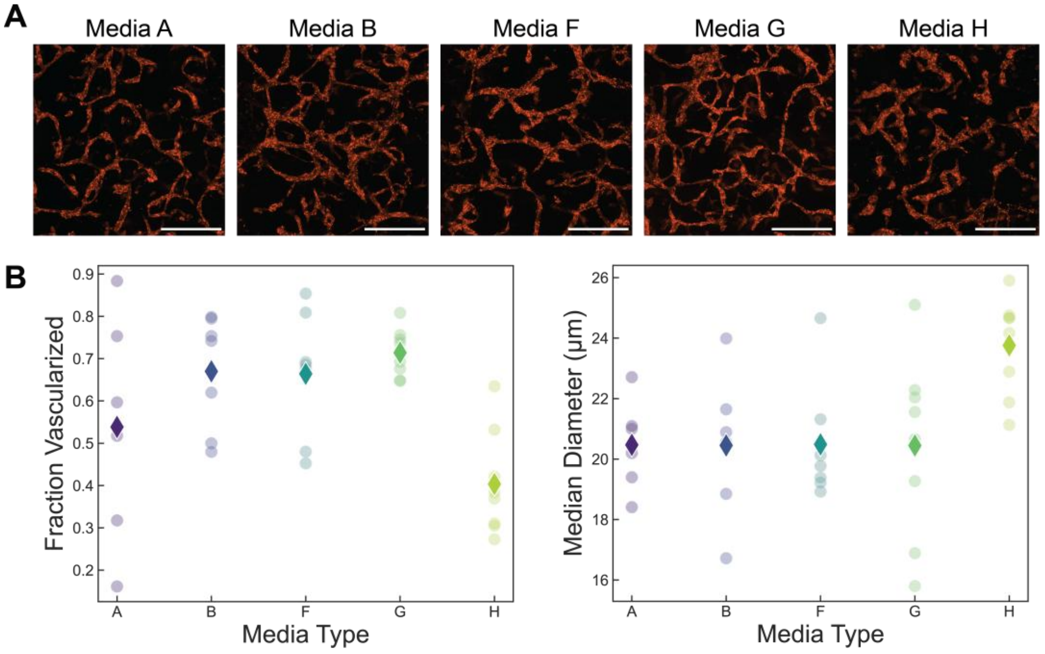


**Fig. S2. Optimization of cell culture medium.** (**A**) Representative images of mCherry-fluorescent BMECs in Brain-Chips in different media formulations as detailed in **Table S1** (day 9; scale bars, 200 μm. (**B**) Corresponding vascular network quantification (n=6-8 chips per group; individual replicates shown as semi-transparent circular markers, group means as solid diamond markers). Fetal bovine serum (FBS) was removed due to predicted neurotoxicity. Initial media screen formulations were guided by the miBrain system.[2]

**Table S1. Media formulation screen for Brain-Chips.** Blue text denotes components of Lonza EGM-2 MV kit, with proprietary concentrations listed as dilutions.

| Media A | Media B | Media F | Media G | Media H | Concentration |
|---|---|---|---|---|---|
| Gentamicin | Gentamicin | Gentamicin | Gentamicin | Gentamicin | 0.1% v/v (kit) |
| | | NEAA | | NEAA | 1% v/v |
| | | CD Lipids | | CD Lipids | 1% v/v |
| | AGS | AGS | AGS | AGS | 1% v/v |
| | B27 | B27 | B27 | B27 | 2% v/v |
| | | | cAMP | cAMP | 1 μM |
| Ascorbic acid | Ascorbic acid | Ascorbic acid | Ascorbic acid | Ascorbic acid | 0.1% v/v (kit) |
| | | | NT3 | NT3 | 10 ng/mL |
| R3-IGF-1 | R3-IGF-1 | R3-IGF-1 | R3-IGF-1 | R3-IGF-1 | 0.1% v/v (kit) |
| FBS | | | | | 5% v/v |
| Hydrocortisone | Hydrocortisone | Hydrocortisone | Hydrocortisone | Hydrocortisone | 0.04% v/v (kit) |
| hFGF-B | hFGF-B | hFGF-B | hFGF-B | hFGF-B | 0.4% v/v (kit) |
| VEGF | VEGF | VEGF | VEGF | VEGF | 0.1% v/v (kit) |
| hEGF | hEGF | hEGF | hEGF | hEGF | 0.1% v/v (kit) |
| SAG | SAG | SAG | SAG | SAG | 1 nM |
| VEGF | VEGF | VEGF | VEGF | VEGF | 50 ng/mL |
| M-CSF | M-CSF | M-CSF | M-CSF | M-CSF | 25 ng/mL |

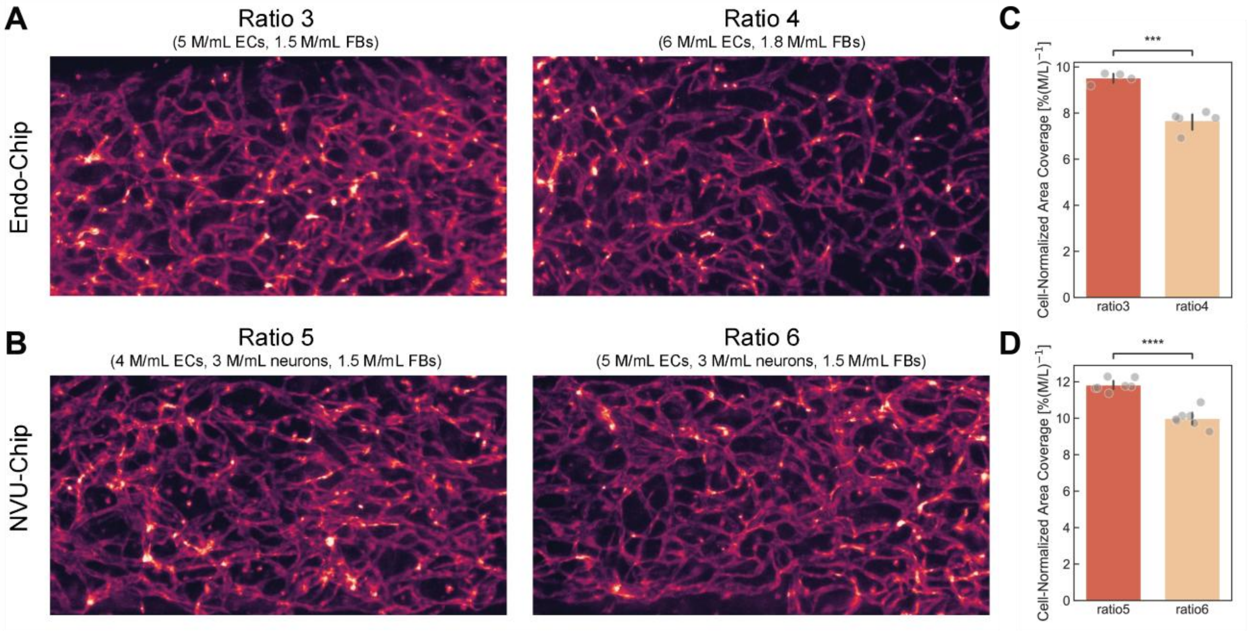


**Fig. S3. Optimization of cell ratios.** Representative images of (**A**) Endo-Chips and (**B**) NVU-Chips at day 8 with two cell different ratios as listed. Corresponding quantification of cell-normalized vascular coverage for (**C**) Endo-Chips and (**D**) NVU-Chips (n=4-7 chips per group). **** $p < 0.0001$, *** $p < 0.001$, ** $p < 0.01$, * $p < 0.05$, ns $p > 0.05$.

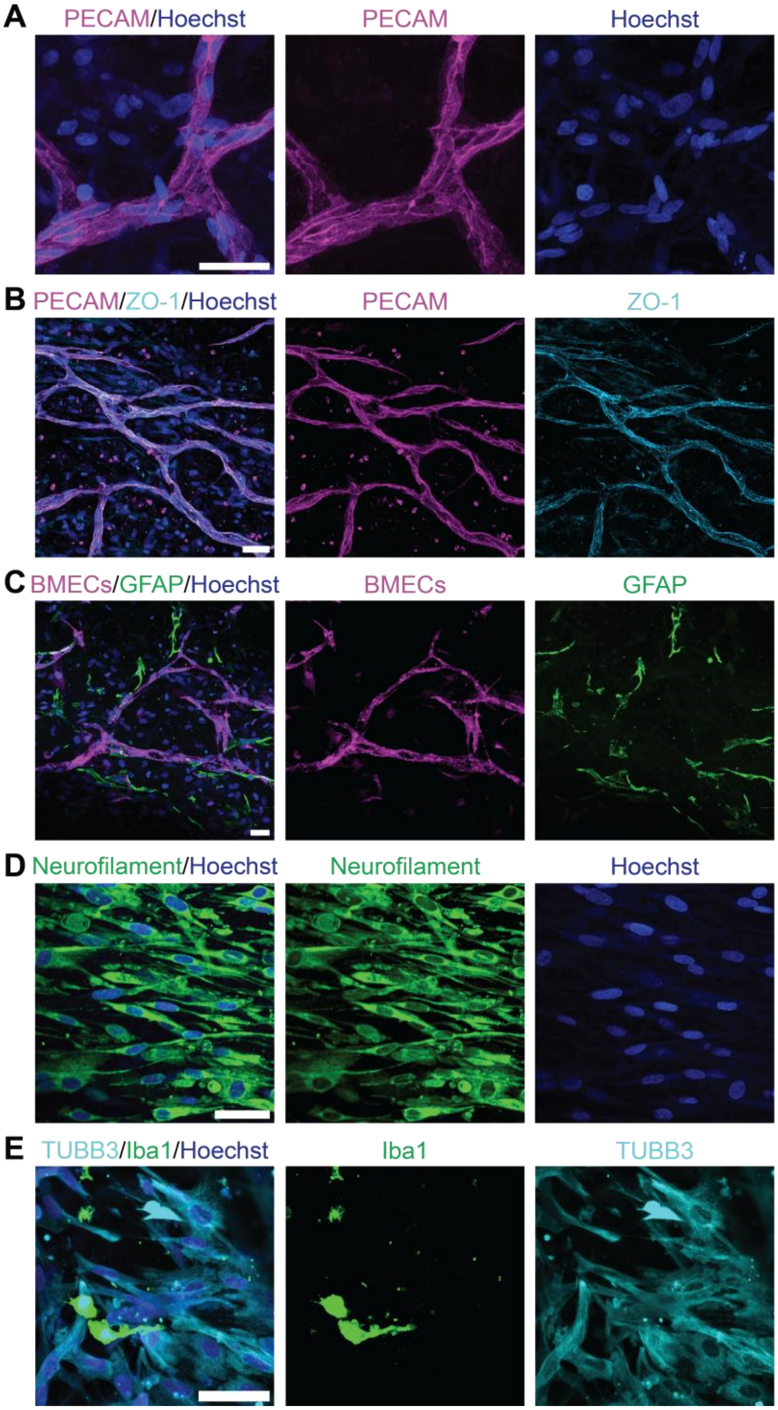


**Fig. S4: Histological characterization of Brain-Chip cultures.** Immunohistochemistry on Brain-Chip cultures for (**A**) BMEC marker PECAM (magenta: PECAM, blue: Hoechst), (**B**) tight junctions (cyan: ZO-1, magenta: PECAM, blue: Hoechst), (**C**) astrocytes (green: GFAP, magenta: mCherry-BMECs, blue: Hoechst), (**D**) neurons (green: neurofilament, blue: Hoechst), and (**E**) iMG (green: Iba1, cyan: TUBB3, blue: Hoechst). All scale bars, 50 µm.

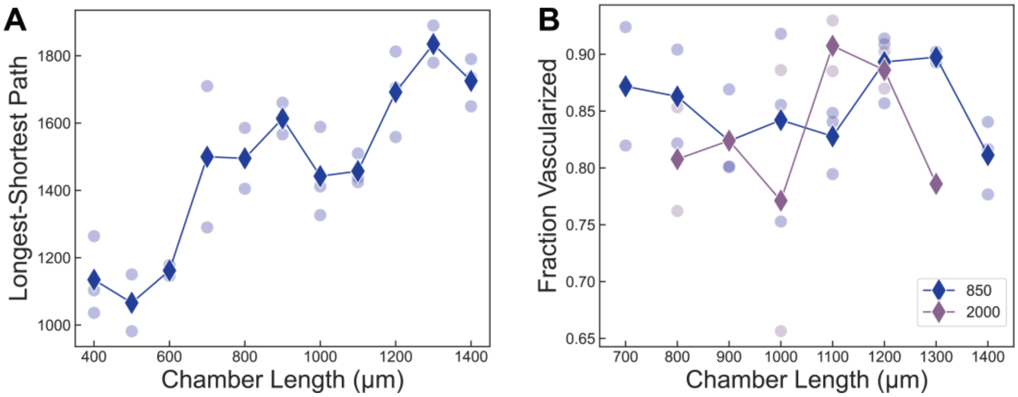


**Fig. S5. Vessel network analysis with varying GelChip design parameters.** (**A**) Longest-shortest path and (**B**) fraction vascularized for Endo-Chips as a function of chamber length (blue: 850 μm chamber width; purple: 2000 μm chamber width; n=2-3 chips per group; individual replicates shown as semi-transparent circular markers, group means as solid diamond markers).

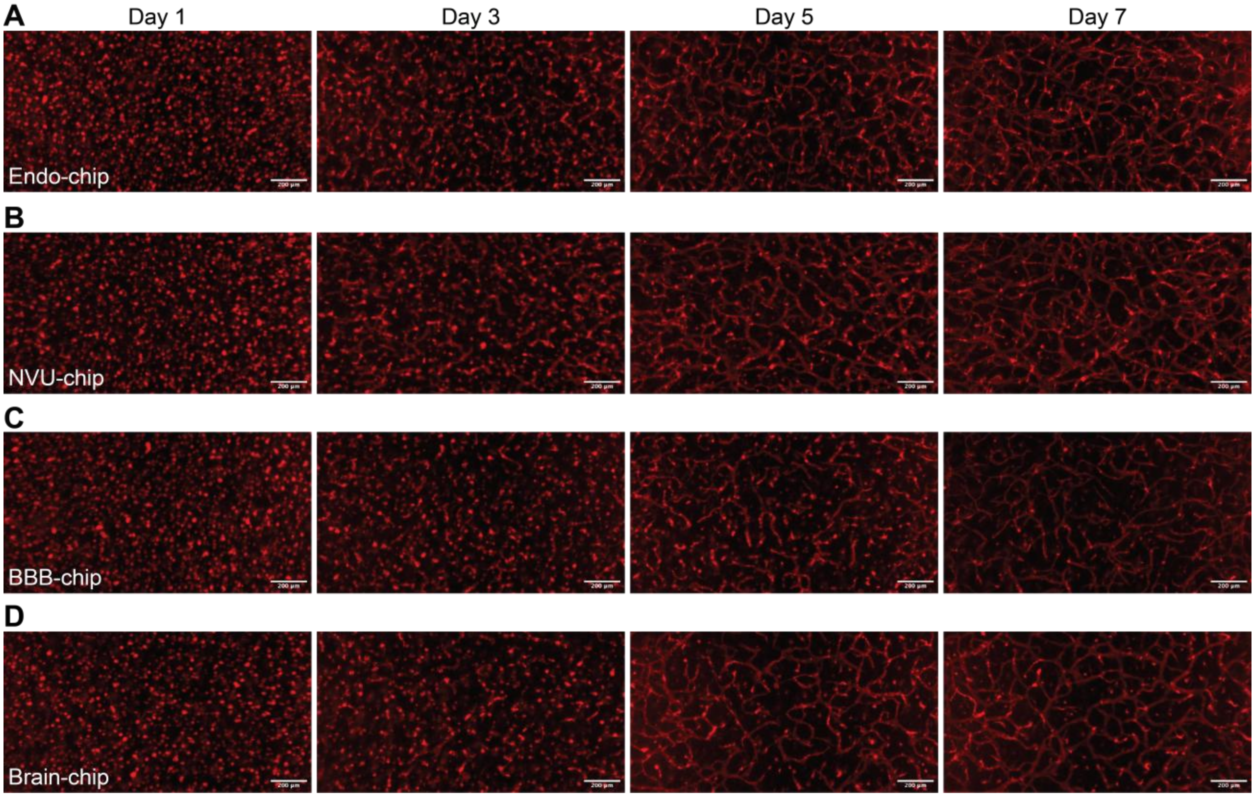


**Fig. S6. Daily vessel images for all chip co-cultures.** BMECs live-imaged as a function of days in culture for (A) Endo-Chip (repeated from main text), (B) NVU-Chip, (C) BBB-Chip, and (D) Brain-Chip (red: mCherry-BMECs; scale bars, 200 μm).

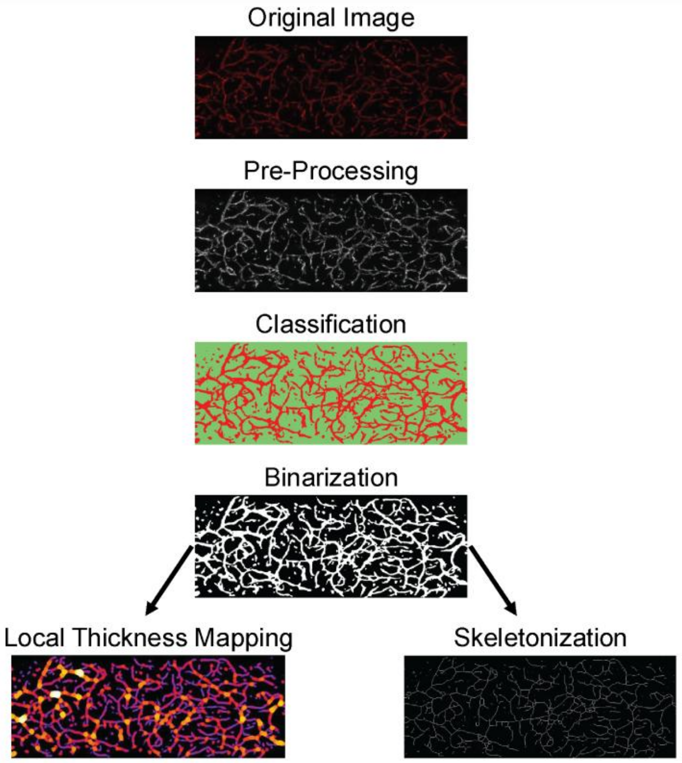


**Fig. S7. Supervised machine-learning–based vessel network analysis.** Workflow as detailed in the Methods: (i) pre-processing, (ii) classification of vessel vs. matrix with Trainable Weka Segmentation, (iii) binarization to a vessel mask, then either: local thickness mapping on the mask with BoneJ or skeletonize and Analyze Skeleton.

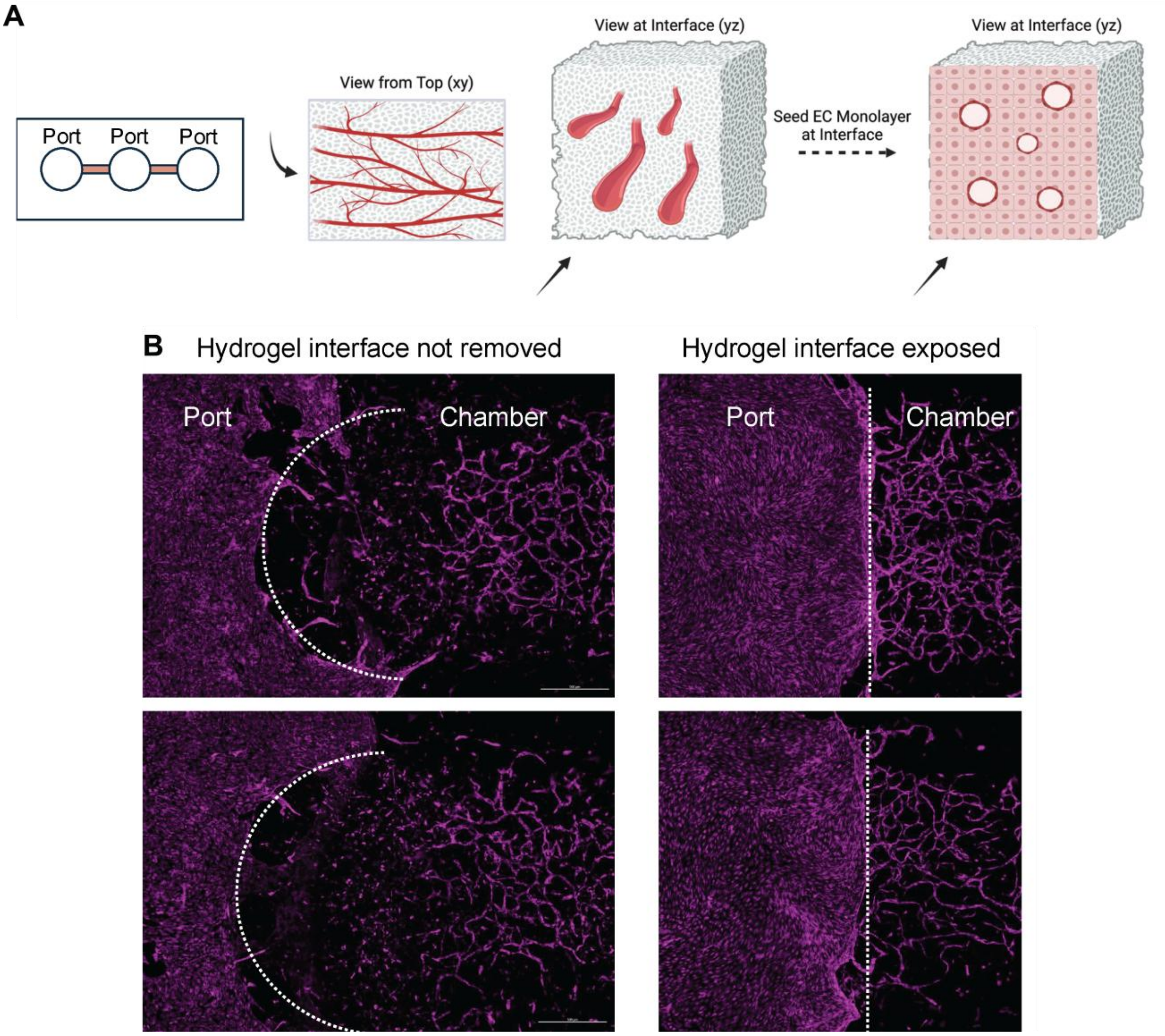


**Fig. S8. Accessing vessel networks in hydrogel via microdissection.** (**A**) Schematic of microfluidic chip interface seeding strategy. (**B**) Representative images depicting (left) hydrogel interface left intact for interface seeding with BMECs vs. (right) hydrogel interface dissected prior to interface seeding (magenta: mCherry-BMECs; scale bars, 500 µm). Dashed white lines added to denote approximate hydrogel edge.
